# Breastfeeding and risk of asthma, hay fever and eczema

**DOI:** 10.1101/144352

**Authors:** Weronica E. Ek, Torgny Karlsson, Carlos Azuaje Hernandez, Mathias Rask-Andersen, Åsa Johansson

**Affiliations:** Department of Immunology, Genetics and Pathology, Science for Life Laboratory, Uppsala University, Uppsala, Sweden

**Keywords:** Breastfeeding, asthma, hay fever, eczema, smoke

## Abstract

**Background:** Breastfeeding is commonly proposed to protect against atopic diseases. However, studies aiming to quantify these protective effects have shown conflicting results.

**Methods:** To entrench the effects of breastfeeding on risk of asthma, hay fever and eczema, our study included a large study cohort, UK Biobank (N=502,682). Information was collected on whether participants had been breastfeed and on the prevalence of disease. Disease was tested for association with breastfeeding, adjusting or matching for influential covariates.

**Findings:** A total of 443,068 participants were included in our analyses of which 71·2% had been breastfed. The prevalence of asthma was 11·4 % and 12·7% in the breastfed and non-breastfed groups, and hay fever or eczema (23·9% and 24·8 % in the two groups respectively. When correcting or matching for potential confounders, we could not see any association between being breastfed and asthma. However, there were increased odds of hay fever and eczema among participants that had been breastfed (P=7·78×10^−6^).

**Interpretation:** This study reports that breastfeeding is associated with increased odds of hay fever and eczema but it show no evidence for breastfeeding being associated with asthma diagnosis.

**Funding:** The Swedish Society for Medical Research (SSMF), the Kjell and Märta Beijers Foundation, Göran Gustafssons Foundation, the Swedish Medical Research Council (Project Number 2015-03327), the Marcus Borgström Foundation, the Åke Wiberg Foundation and the Vleugels Foundation.

**Evidence before this study:** Atopic diseases affect quality of life for a large part of the human population and pose a very high socio-economic burden. Genetic, environmental, and a number of lifestyle factors influence our risk of developing atopic disorders and high familial prevalence is one of the strongest known risk factors for disease. Several environmental and lifestyle risk factors have already been well established in the scientific community, such as smoking on the risk of developing asthma. Breastfeeding is commonly argued to be protective against atopic diseases. However, studies aiming to quantify these protective effects have shown conflicting results.

**Added value of this study:** Our study is, to our knowledge, the largest investigation on how breastfeeding is associated with being diagnosed with asthma, hay fever and eczema at adult age. The study found breastfeeding to be associated with increased odds of being diagnosed with hay fever and eczema during life, while we found no association between breastfeeding and asthma. Our results for hay fever and eczema is in line with the western world hygiene hypothesis, but contradict the general picture of breastfeeding being protective.

**Implications of all the available evidence:** To be able to give parents correct advice on lifestyles choices that will protect their kids against atopic diseases, we need to clarify the currently conflicting results on the effect of breastfeeding on risk of atopic diseases. However, these results should not be used to recommend breastfeeding or to discourage it since the present study only investigates the association between breastfeeding history and being diagnosed with asthma, hay fever and eczema during lifetime.

**Abbreviations:**

TDI: Townsend Deprivation Index
BMI: Body Mass Index

## Introduction

According to the World Health Organization’s (WHO) guidelines, breastfeeding is advocated for at least six months after the baby is born because of its associated health benefits for the child^1^. Human milk is the biological norm for feeding newborns and breastfeeding is recommended for primary prevention of atopic diseases^2^. Atopy refers to a tendency to produce immunoglobulin E (IgE) antibodies in response to common environmental proteins such as food allergens, pollen and house dust mite. Individuals with a presence of atopy also have an increased risk of developing atopic diseases, such as asthma, hay fever and eczema. Atopic diseases affect quality of life for a large part of the human population and pose a very high socio-economic burden^3^. Patients often require lifetime treatment, and in severe cases even hospitalization. Asthma and allergies can also be life threatening, and despite the availability of a large number of medications for symptomatic relief, at present, no curative treatment is available. Genetic, environmental, and a number of lifestyle factors influence our risk of developing atopic disorders and high familial prevalence is one of the strongest known risk factors for disease. Several environmental and lifestyle risk factors have already been well established in the scientific community, such as smoking on the risk of developing asthma^4^. Other factors, such as breastfeeding, have yielded more inconsistent results, on the risk of developing asthma, hay fever and eczema^5,6,7^.

The view that breastfeeding reduces the risk for atopic disease in the child is widely accepted and promoted. However, published research show conflicting results on the effect of breastfeeding on atopic diseases. Some investigators have reported on a protective effect of breastfeeding^8,9,10^ while others find no effect, or even an increased risk of asthma and eczema in breastfed babies^11,12,13^. Studies have been done on observational data due to ethical constraints; it is not ethically defencible to withdraw breastfeeding from children in a clinical trial. Methodological shortcomings such as the failure to consider important confounders may be one reason for heterogeneity of the results. Other reasons may be differences in the age of participants, publication bias towards positive results, variation in exposures to childhood-infections or differences in socioeconomic status between breastfed and non-breastfed individuals. A general shortcoming with many previous studies is the relatively small sample sizes (N < 2,000), which increase the rate of false negative results (i.e., the type II error) due to low statistical power. To be able to give parents correct advice on lifestyles choices that will protect their children against atopic diseases, we need to clarify the currently conflicting results on the effect of breastfeeding on risk of atopic diseases.

Although breastfeeding can be recommended for many different health benefits, the present study aims to specifically investigate whether breastfeeding alters the longterm risks of asthma, hay fever and eczema. A large homogenous population based cohort, as the one used in this study (UK Biobank N=502,682), provides statistical power to fill in currently unaddressed, yet highly important, gaps in the knowledge on the effects of breastfeeding on the risks of developing asthma, hay fever and eczema later in life.

## Material and Methods

### Participants

The UK Biobank database includes 502,682 participants recruited from all across the UK. Participants were aged 37 to 73 years at the time of recruitment between 2006 and 2010. Most participants visited the center once, but some individuals visited the center at up to three instances. This population-based cohort includes information on medical conditions, diet and lifestyle factors. The UK Biobank database was approved by the National Research Ethics Committee (REC reference 11/NW/0382). Applications for using data from UK Biobank has been approved (application nr: 15479). In this study, we only included participants of Caucasian ancestry, defined as those reporting that they were “white”, “white British”, “Irish” or “other white” during all instances. 5,568 individuals were removed due to conflicting answers on asthma (622 individuals), hay fever (2,143 individuals), and eczema (2,738 individuals) and 65 individuals due to conflicting answers on both hay fever and eczema.

### Disease phenotypes: asthma and hay fever/eczema

Self-reported asthma and hay fever and/or eczema (combined) have been collected through touch screen questionnaire. The primary data used in this study were from the touch screen question number 6152: “*has a doctor ever told you that you have had any of the following conditions? (You can select more than one answer)*”; 1) asthma and 2) hay fever, allergic rhinitis or eczema, 3) none of the above or 4) prefer not to answer. Hay fever and eczema diagnosis could not be separated and this variable is therefore called hay fever/eczema in our study. Controls were selected as individuals answering “none of the above” in question 6152, and those who did not report asthma, hay fever or eczema during the verbal interview (see below). Some individuals visited the center more than once and we included participants that reported disease at any or all visits.

### Disease phenotypes: hay fever and eczema separately

Trained staff followed up individuals who answered that had been told by a doctor that they had one or more of the following diseases: heart attack, angina, stroke, high blood pressure, blood clot in leg, blood clot in lung, emphysema/chronic bronchitis, asthma or diabetes, with a verbal interview. During this interview the participants where shown a tree structure based on ICD-10 classifications and asked to more specifically highlight which diseases they had, including atopic diseases such as eczema or hay fever. For each illness entered, the interviewer recorded the date of the first diagnosis. These diagnoses (UK biobank data field 20002) were used to define hay fever and eczema cases separately. Since only 379,887 participants were followed up for an interview, and this selection was not random, diseases reported here have a bias in prevalence. In addition, diseases in this variable are highly underreported, since the tree structure with ICD10 classifications is huge, and participants were guided into what part of the tree to look at. Therefore, data from individuals who did not report that they had eczema or hay fever is uncertain and the same controls were therefore used as for the touchscreen variables above. If there were any conflicting answers between the variables from the interview and the touchscreen, the phenotype was set to missing. For example, individuals who reported that they had eczema/hay fever in the touch screen questionnaire, but had not reported either eczema or hay fever in the interview were removed from the analyses.

### Statistical analysis

Statistic analysis was performed using the stats library of R version 2.15.10^14^. All disease phenotypes were coded as binary (1 = yes, and 0 = no), and generalized linear models with a logit link-function, i.e., logistic regression, were applied when calculating odds ratios. In the first analysis, we compared the odds of the diseases depending on breastfeeding without including any covariates. We then re-analyzed all disease phenotypes and included the covariates that we identified to be associated to the disease phenotypes or breastfeeding in our study (Table 2). To measure the robustness of the associations, we did additional subgroup analyses separately for men and women, being exposed to smoke or not and if the participants where born during or before 1950 or after 1950. Stratifying for being born before or after 1950 were done since year of birth was highly significant to both the chance of being breastfed and to disease risk. After 1950 there was a quicker increase in disease incidence and at the same time point a fast decline in the incidence of being breastfed (Figure 1).

**Figure 1.**
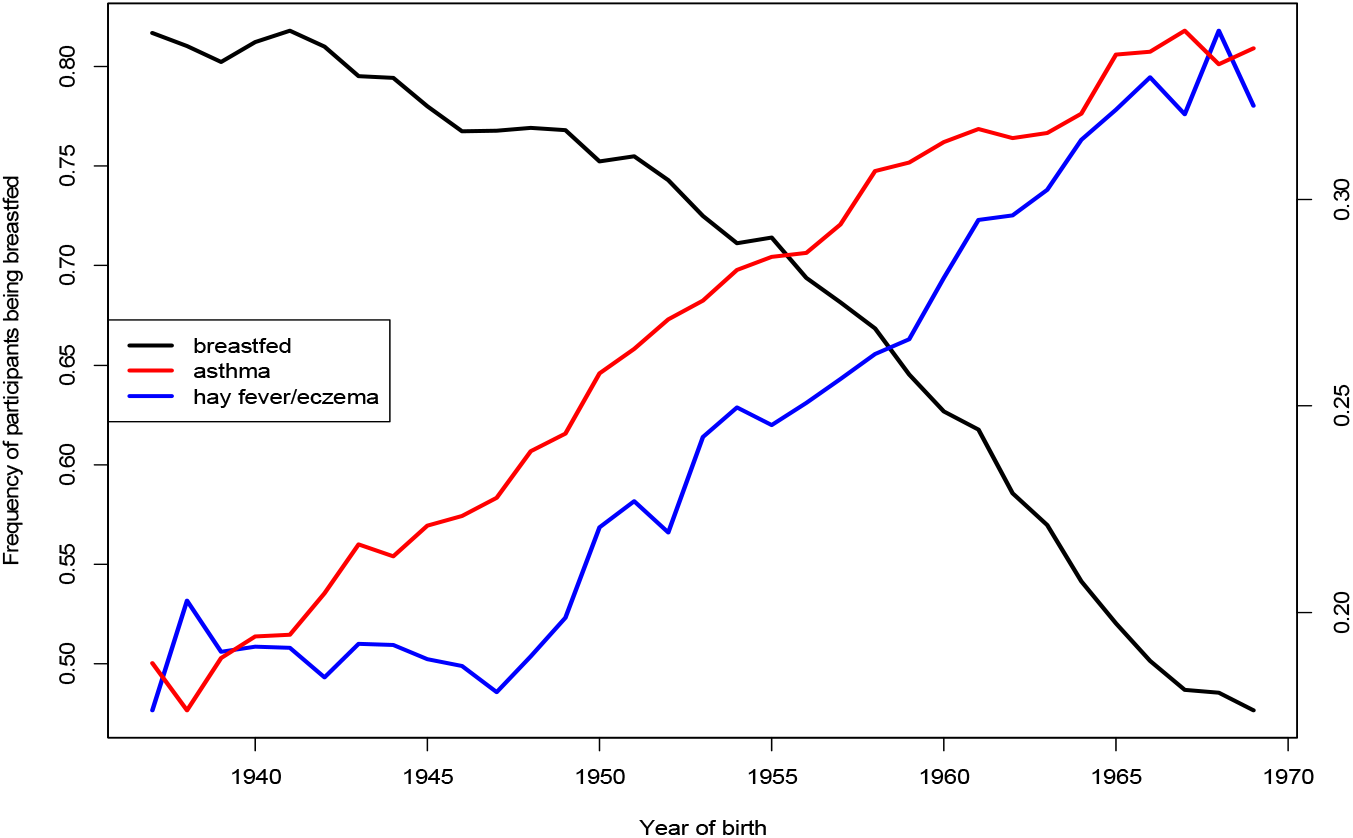
Frequency of participants being breastfeed and the incidence of asthma, hay fever/eczema for year of birth from 1934 to 1970.

#### Matching breastfed and non-breastfed participants

To further strengthen the results from our main analysis and minimize loss of power due to model misspecification, we further matched breastfed and non-breastfed participants for possible confounders. First, participants with a birth weight lower than 2·5 kg (N=25,612) and participants with a BMI lower than 18·5 (N= 2,266) or higher than 29·9 (N=109,881) were removed prior to analysis. Participants were thereafter stratified into being born before or after 1950, males or females, having a Townsend deprivation index (TDI) below or over the median value in our sample (median=−2·44), exposed or not exposed to smoke (either by ever smoked themselves, living with someone who smokes or having a mother that smoked around birth). This created 16 groups with matched breastfed and non-breastfed participants (Figure 2). We compared the odds of disease on breastfeeding for each matched group, adjusting for year of birth, birth weight, BMI and TDI. These 16 groups where thereafter metaanalyzed using both a fixed and a random effects model.

**Figure 2.**
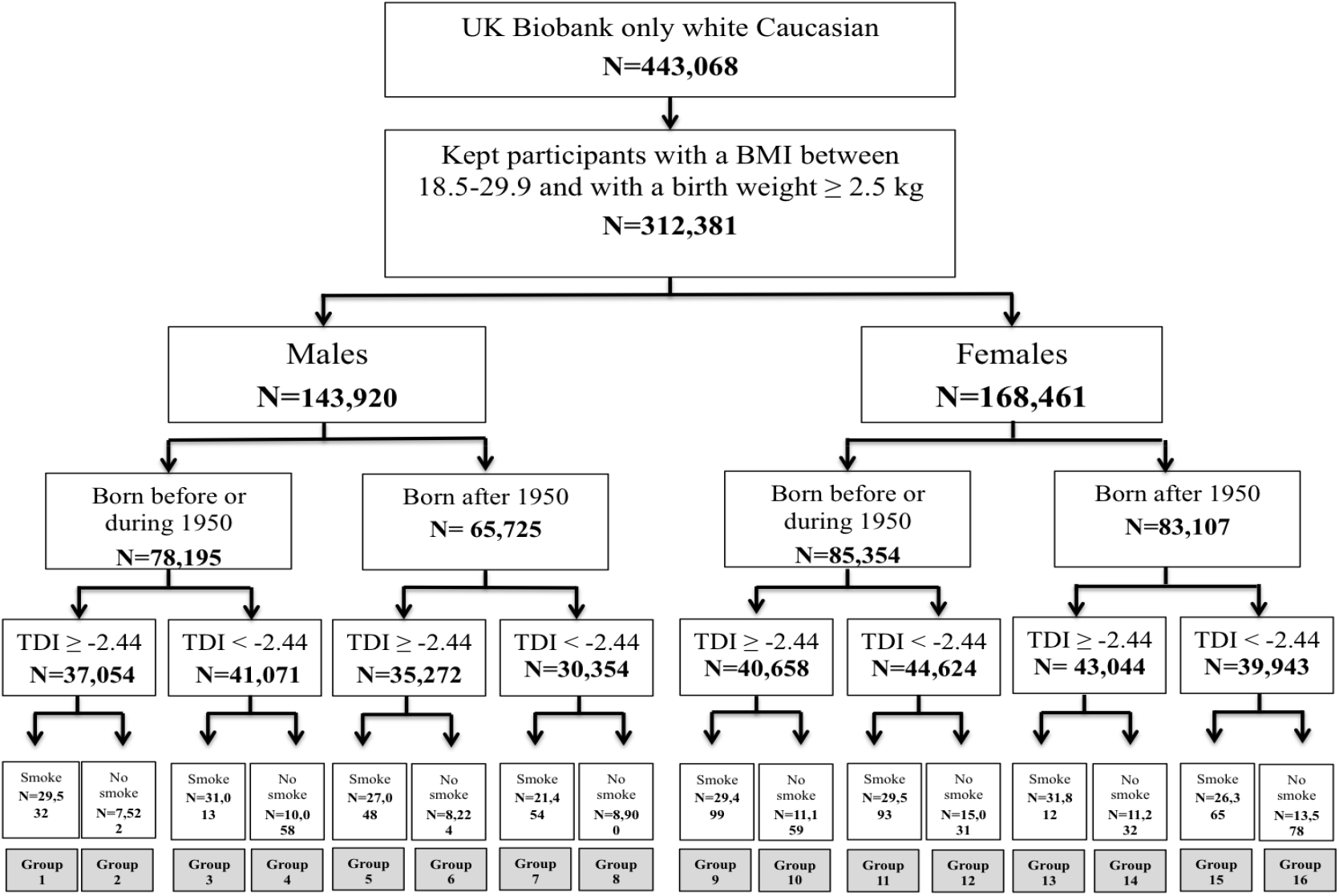
Flow chart of matched groups.

## Results

A total of 443,068 Caucasian participants were included in this study, of which 106,704 participants did not provide an answer to whether they were breastfed or not, or answered that they did not know. Among the remaining 336,364, 239,239 (71 ·2 %) answered yes to having been breastfed, and 97,125 (28·8 %) answered no. Baseline characteristics and prevalence for each group are presented in Table 1. The number of cases for the disease phenotypes was: 51,645 for asthma, 102,862 for hay fever/eczema, 22,919 for hay fever and 9,578 for eczema. A total of 294,477 participants were selected as healthy controls. Frequency of being breastfed decreased dramatically with year born form >80% in participants born between 1937 and 1941 to lower than 50 % during 1970 (Figure 2). Oppositely, incidence of asthma increased dramatically in participants born between 1935 and 1970 and hay fever/eczema increased quickly in participants born between 1950 and 1971 (Figure 1).

**Table 1.**
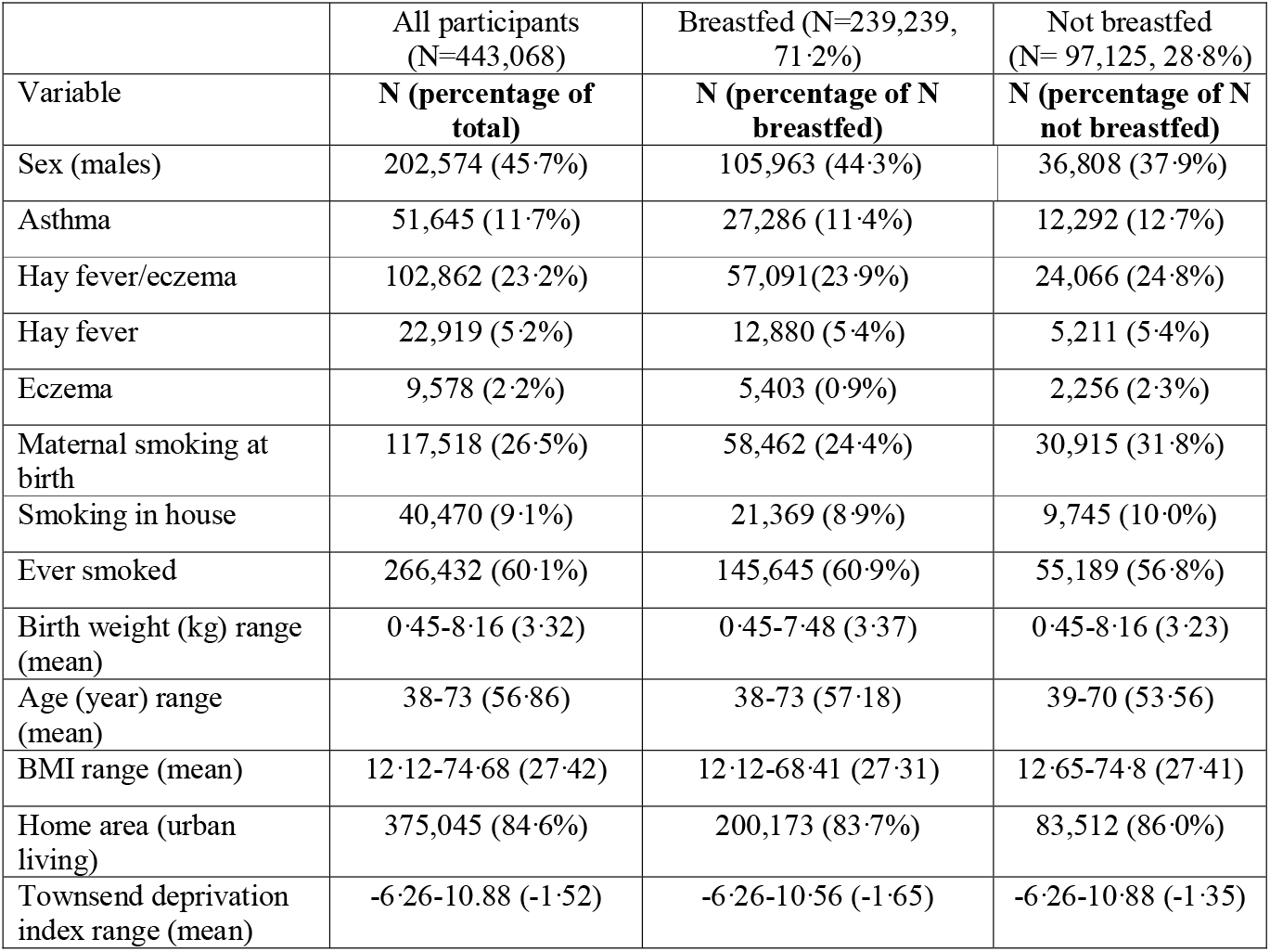
Baseline characteristics for the participants include in the study.

|  | All participants<br>(N=443,068) | Breastfed (N=239,239,<br>71.2%) | Not breastfed<br>(N= 97,125, 28.8%) |
| --- | --- | --- | --- |
| Variable | **N (percentage of**<br>**total)** | **N (percentage of N**<br>**breastfed)** | **N (percentage of N**<br>**not breastfed)** |
| Sex (males) | 202,574 (45.7%) | 105,963 (44.3%) | 36,808 (37.9%) |
| Asthma | 51,645 (11.7%) | 27,286 (11.4%) | 12,292 (12.7%) |
| Hay fever/eczema | 102,862 (23.2%) | 57,091 (23.9%) | 24,066 (24.8%) |
| Hay fever | 22,919 (5.2%) | 12,880 (5.4%) | 5,211 (5.4%) |
| Eczema | 9,578 (2.2%) | 5,403 (0.9%) | 2,256 (2.3%) |
| Maternal smoking at<br>birth | 117,518 (26.5%) | 58,462 (24.4%) | 30,915 (31.8%) |
| Smoking in house | 40,470 (9.1%) | 21,369 (8.9%) | 9,745 (10.0%) |
| Ever smoked | 266,432 (60.1%) | 145,645 (60.9%) | 55,189 (56.8%) |
| Birth weight (kg) range<br>(mean) | 0.45-8.16 (3.32) | 0.45-7.48 (3.37) | 0.45-8.16 (3.23) |
| Age (year) range<br>(mean) | 38-73 (56.86) | 38-73 (57.18) | 39-70 (53.56) |
| BMI range (mean) | 12.12-74.68 (27.42) | 12.12-68.41 (27.31) | 12.65-74.8 (27.41) |
| Home area (urban<br>living) | 375,045 (84.6%) | 200,173 (83.7%) | 83,512 (86.0%) |
| Townsend deprivation<br>index range (mean) | -6.26-10.88 (-1.52) | -6.26-10.56 (-1.65) | -6.26-10.88 (-1.35) |

### Association between disease and individual covariates

We did see a strong association between the diseases and most of the tested covariates (Table 2). TDI was associated with both asthma and hay fever/eczema. For asthma, higher TDI (i.e. lower income and lower education level) was associated with larger odds of being diagnosed with asthma and lower odds of being diagnosed with hay fever or eczema. Maternal smoking around birth and smoking in the house seemed to be risk factors for developing asthma, while it had a protective effect on being diagnosed with hay fever and eczema. If participants had ever smoked was not associated with asthma diagnosis (P=0·91), hay fever (P=0·57) or eczema (P=0·33) but showed a significant protective effect for hay fever/eczema OR=0·92). Home area (urban or rural living) was not significantly associated with hay fever /eczema (P=0·12) but rural living lowered the odds of hay fever and eczema independently (OR=0·82 and OR=0·91, respectively) and increased the odds of asthma (OR=1·03, Table 2). A higher BMI increased the odds of being diagnosed with asthma (OR=1·04), while it lowered the risk of being diagnosed with hay fever or eczema (OR=0·99).

**Table 2.** Associations between individual covariates and asthma, hay fever, eczema and breastfeeding.

|  | Asthma |  | Hay fever<br>/eczema |  | Hay fever |  | Eczema |  | Breastfeeding |  |
| --- | --- | --- | --- | --- | --- | --- | --- | --- | --- | --- |
|  | **OR**<br>**(95 % CI)** | **P** | **OR**<br>**(95 % CI)** | **P** | **OR**<br>**(95 % CI)** | **P** | **OR**<br>**(95 % CI)** | **P** | **OR**<br>**(95 % CI)** | **P** |
| Sex | 0.816<br>(0.801-0.832) | <2x10<sup>-16</sup> | 0.796<br>(0.784-0.807) | <2x10<sup>-16</sup> | 0.955<br>(0.930-0.981) | 0.0009 | 0.837<br>(0.803-0.872) | <2x10<sup>-16</sup> | 1.302<br>(1.283-1.323) | <2x10<sup>-16</sup> |
| TDI* | 1.029<br>(1.026-1.032) | <2x10<sup>-16</sup> | 0.990<br>(0.987-0.992) | <2x10<sup>-16</sup> | 0.960<br>(0.956-0.965) | <2x10<sup>-16</sup> | 0.993<br>(0.986-1.000) | 0.038 | 0.966<br>(0.964-0.968) | <2x10<sup>-16</sup> |
| Birth weight | 0.918<br>(0.901-0.935) | <2x10<sup>-16</sup> | 0.974<br>(0.961-0.988) | 0.0002 | 0.978<br>(0.952-1.003) | 0.087 | 0.994<br>(0.956-1.034) | 0.78 | 1.401<br>(1.381-1.421) | <2x10<sup>-16</sup> |
| Year of birth | 1.012<br>(1.016-1.019) | <2x10<sup>-16</sup> | 1.030<br>(1.029-1.031) | <2x10<sup>-16</sup> | 1.033<br>(1.032-1.035) | <2x10<sup>-16</sup> | 1.034<br>(1.031-1.036) | <2x10<sup>-16</sup> | 0.946<br>(0.945-0.947) | <2x10<sup>-16</sup> |
| Maternal<br>Smoke | 1.123<br>(1.099-1.147) | <2x10<sup>-16</sup> | 0.962<br>(0.946-0.978) | 4x10<sup>-6</sup> | 0.902<br>(0.874-0.932) | 3x10<sup>-10</sup> | 0.940<br>(0.896-0.986) | 0.011 | 0.665<br>(0.654-0.676) | <2x10<sup>-16</sup> |
| Smoke in<br>house | 1.107<br>(1.073-1.142) | <2x10<sup>-16</sup> | 0.972<br>(0.948-0.997) | 0.027 | 0.920<br>(0.878-0.965) | 0.0006 | 0.947<br>(0.881-1.018) | 0.141 | 0.864<br>(0.842-0.886) | <2x10<sup>-16</sup> |
| Ever smoked | 0.998<br>(0.978-1.017) | 0.91 | 0.923<br>(0.910-0.937) | <2x10<sup>-16</sup> | 0.990<br>(0.958-1.024) | 0.57 | 1.021<br>(0.979-1.065) | 0.33 | 1.184<br>(1.166-1.202) | <2x10<sup>-16</sup> |
| Home area** | 1.031<br>(1.004-1.059) | 0.025 | 0.984<br>(0.964-1.004) | 0.122 | 0.818<br>(0.789-0.848) | <2x10<sup>-16</sup> | 0.912<br>(0.86-0.95) | 0.001 | 0.969<br>(0.966-0.972) | <2x10<sup>-16</sup> |
| BMI | 1.035<br>(1.033-1.037) | <2x10<sup>-16</sup> | 0.996<br>(0.995-0.998) | 4x10<sup>-7</sup> | 0.99<br>(0.988-0.99) | 10x10<sup>-8</sup> | 0.996<br>(0.993-1.000) | 0.035 | 0.992<br>(0.988-0.995) | 9x10<sup>-7</sup> |

\*TDI = Townsend Deprivation Index
\*\*Home area is either rural or urban

### Association between breastfeeding and individual covariates

Breastfeeding, the predictor variable of interest, was highly associated with all covariates tested (Table 2). The odds of being breastfed were lower in individuals with high TDI (i.e. individuals with lower socioeconomic status, OR=0·95), in those who were born later (OR=0·82), and in those whose mother smoked around birth (OR=0·92) (Table 2). There was no strong evidence of multicollinearity between breastfeeding and the covariates, as suggested by the low variance inflation factor associated with breastfeeding (VIF_breastfeeding_ = 1·06). Therefore, the potential problem of variance and misspecification bias inflation may safely be disregarded.

### Unadjusted results

Without considering covariates, we observe breastfeeding to have a protective effect on both asthma (OR=0·88, P<2.10×10^−16^, Figure 3A) and hay fever/eczema (OR=0·94, P=1·89×10^−10^, Figure 3B). However, when eczema and hay fever were diagnosed separately (using variable 20002), we could not see any statistically significant effect on either eczema (OR=0·98, P=0·34) or hay fever (OR=0·95, P=0·060) (Figure 3C and 3D).

**Figure 3 A-D.**
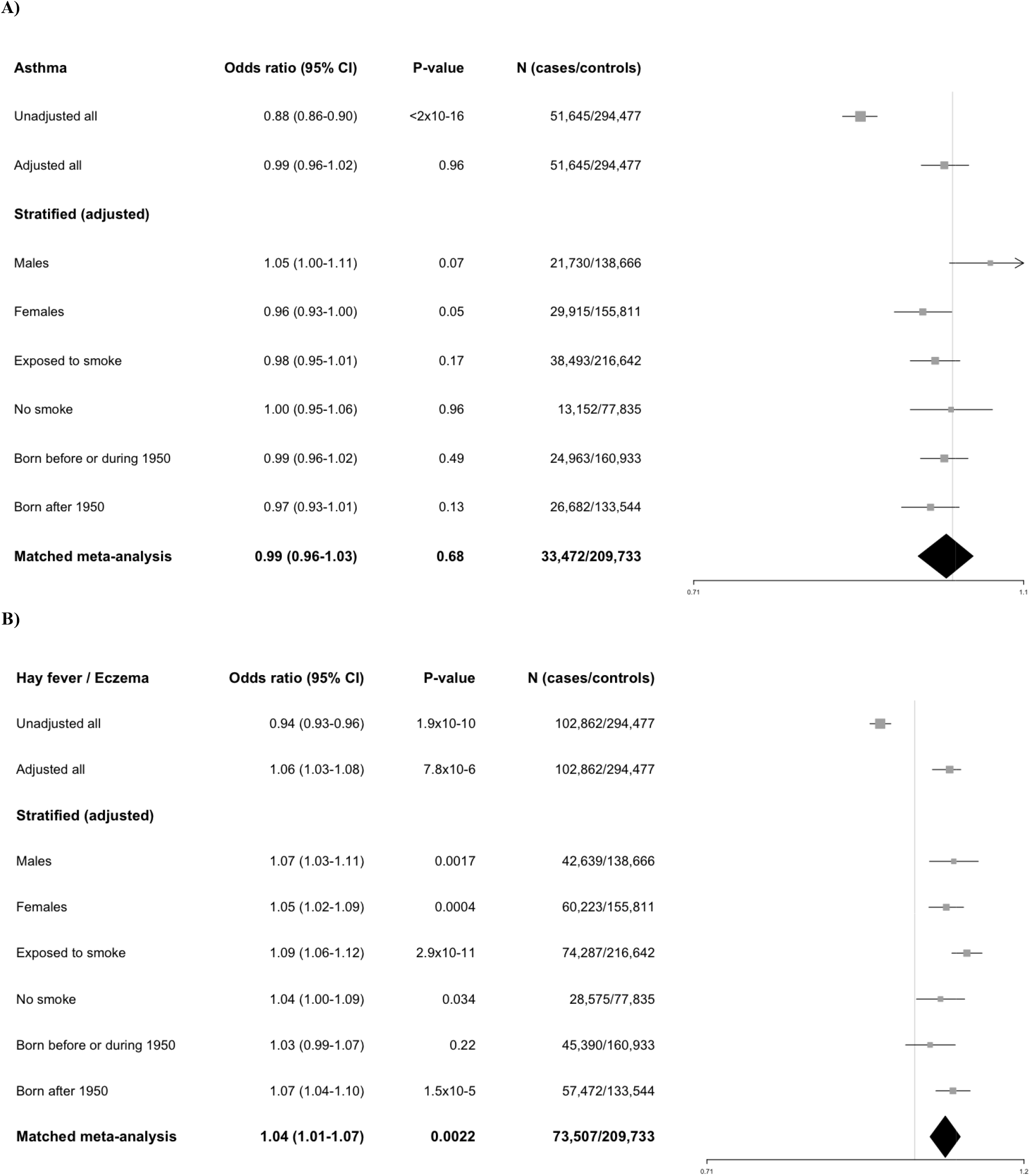
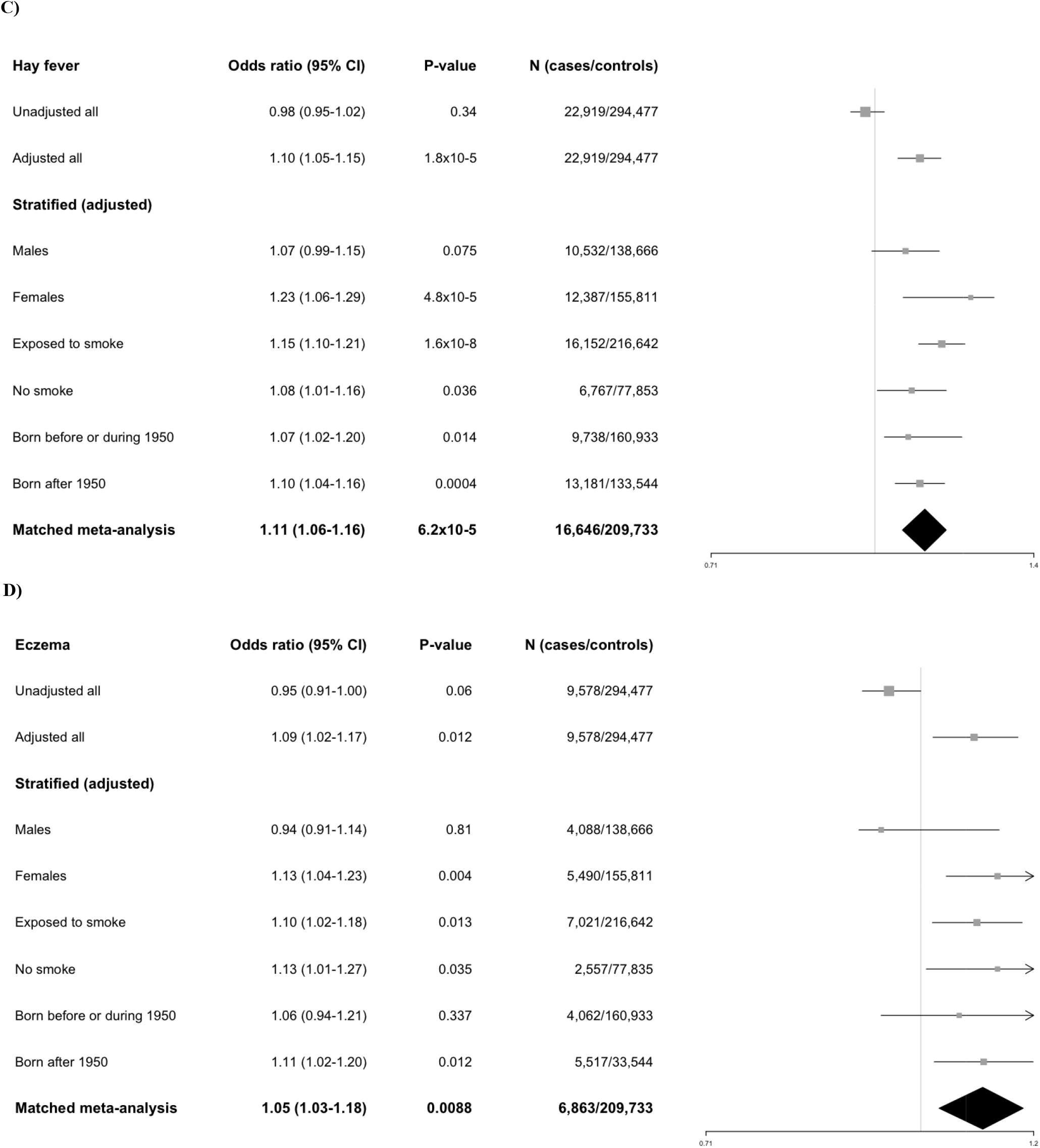
Forest plots for the effect of breastfeeding on asthma and hay fever/eczema for the unadjusted model, the adjusted model, the adjusted stratified analysis (stratified for sex, smoke exposure and year of birth) and the matched meta-analysis for **A**) Asthma, **B**) Hay fever / Eczema **C**) Hay fever and D) Eczema. The adjusted model included year of birth, sex, BMI, home area, birth weight, ever smoked, maternal smoking around birth, smoking in house and Townsend deprivation index as covariates in the model. The matched meta-analysis is adjusted for BMI, birth weight, TDI, home area and year of birth.

### Adjusted results

In order to evaluate the effect of the covariates, each covariate was first included in the model separately (Table 3). By including year of birth in the model we can clearly see that the effect of breastfeeding changes from having a protective effect on the odds of being diagnosed with hay fever and/or eczema, to increasing the odds of disease. This suggests that year of birth is a strong confounder in the unadjusted analyses (cf. Figure 2).

**Table 3.** Association between breastfeeding and each disease when including a single covariate.

|  | Asthma |  | Hay fever / eczema |  | Hay fever |  | Eczema |  |
| --- | --- | --- | --- | --- | --- | --- | --- | --- |
| Included covariate | OR (95%CI) | P | OR (95%CI) | P | OR (95%CI) | P | OR (95%CI) | P |
| Sex | 0.893 (0.873-0.914) | $<2 \times 10^{-16}$ | 0.957(0.940-0.974) | $9 \times 10^{-7}$ | 0.985 (0.953-1.019) | 0.379 | 0.963 (0.916-1.012) | 0.138 |
| TDI | 0.891 (0.870-0.912) | $<2 \times 10^{-16}$ | 0.942 (0.925-0.959) | $3 \times 10^{-11}$ | 0.975 (0.942-1.008) | 0.133 | 0.951 (0.904-1.000) | 0.048 |
| Birth weight | 0.918 (0.893-0.944) | $2 \times 10^{-10}$ | 0.977 (0.956-0.997) | 0.026 | 1.007 (0.968-1.048) | 0.713 | 0.970 (0.913-1.029) | 0.310 |
| Year of birth | 0.935 (0.913-0.957) | $3 \times 10^{-8}$ | 1.047 (1.028-1.066) | $8 \times 10^{-8}$ | 1.113 (1.076-1.152) | $10 \times 10^{-10}$ | 1.080 (1.026-1.137) | 0.003 |
| Maternal Smoke | 0.899 (0.877-0.921) | $<2 \times 10^{-16}$ | 0.938 (0.921-0.956) | $3 \times 10^{-10}$ | 0.982 (0.947-1.017) | 0.309 | 0.948 (0.899-1.000) | 0.051 |
| Smoke in house | 0.888 (0.866-0.909) | $<2 \times 10^{-16}$ | 0.940 (0.922-0.957) | $3 \times 10^{-11}$ | 0.971 (0.938-1.005) | 0.093 | 0.949 (0.901-0.999) | 0.047 |
| Ever smoked | 0.883 (0.863-0.904) | $<2 \times 10^{-16}$ | 0.948 (0.931-0.964) | $2 \times 10^{-9}$ | 0.990 (0.958-1.024) | 0.566 | 0.955 (0.908-1.004) | 0.069 |
| BMI | 0.888 (0.868-0.909) | $<2 \times 10^{-16}$ | 0.944 (0.927-0.961) | $1 \times 10^{-10}$ | 0.984 (0.952-1.018) | 0.345 | 0.952 (0.905-1.001) | 0.053 |
| Home area | 0.884 (0.905-0.864) | $<2 \times 10^{-16}$ | 0.944 (0.927-0.961) | $2 \times 10^{-10}$ | 0.978 (0.946-1.012) | 0.195 | 0.948 (0.902-0.997) | 0.021 |

When adjusting for all considered covariates in the full model (i.e. TDI, BMI, rural or urban living, sex, year of birth, birth weight, maternal smoking around birth, smoking in house and if the participant ever smoked), the effect of breastfeeding on the odds of being diagnosed with asthma was no longer significant (OR=0·99, P=0·53, Figure 3A). Hay fever and eczema diagnosed together (hay fever/eczema, variable 6152) was associated with being breastfed, now showing a positive association, i.e. being breastfed as a baby is associated with a higher risk of being diagnosed with disease (OR=1·06, P=2·87×10^−6^) (Figure 2B). Breastfeeding also increase the odds for eczema and hay fever diagnosed separately (variable 20002) (OR=1·09, P=0·012; OR=1·1·01, P=1·77×10^−5^, respectively) (Figure 3C-D).

The adjusted full model also showed that maternal smoking around birth is significantly associated to increased odds of asthma (OR=1·03 (1·00-1·07 95% CI), P=0·047) and lower odds of being diagnosed with hay fever /eczema (variable 6152) (OR=0·94 (0·92-0·97 95% CI), P=3·54×10^−6^). High TDI (i.e. lower income and education) was significantly associated with increased odds of asthma (OR=1·02 (1·01-1·02 95 % CI), P=9·01×10^−11^), but was not statistically associated with hay fever / eczema (OR=0·996, P=0·063). High BMI was significantly associated with increased odds of asthma (OR=1·04 (1·03-1·04 95%CI), P=<2×10^−16^) and hay fever / eczema (OR=1·004 (1·00-1.01 95% CI), P=0·00024), but higher birth weight lowered the odds of asthma (OR=0·93 (0·90-0·95 95% CI), P=3·13×10^−11^) and hay fever / eczema (OR=0·97 (0·96-0·99 95% CI), P=0·0039).

### Stratified analysis

We then stratified for one variable at the time. For the unadjusted analysis, all stratified groups showed a significant protective effect of breastfeeding on asthma, similar as to the unadjusted model including all participants (Supplementary Table 1). However, only females showed a significant association between breastfeeding and asthma in the adjusted analysis (P=0·046), with an opposite effect compared to males (Supplementary Table 1 and Figure 3 A). This result would however not hold for multiple testing. Breastfeeding showed a significant increased odds of hay fever/eczema in participants being born after 1950, but not in participants born during or before 1950 for both the unadjusted and adjusted model stratified for year born (Supplementary Table 1 and Figure 3 B). The odds where similar between all other stratified groups (Supplementary Table 1 and Figure 3 A-D).

### Matching breastfed and non-breastfed participants

After matching for possible confounders, a total of 312,011 Caucasian participants with a BMI between 18·5-29·9 and with a birth weight ≥ 2·5 kg were included in the analysis. Each matched group is defined in Figure 2 and the association results for each group is presented in Supplementary Table 2. When meta-analyzing the 16 groups, breastfeeding was still significantly associated with an increased risk of being diagnosed with hay fever/eczema, hay fever and eczema (Table 4). These results are similar as to the adjusted full model and further suggest that these associations are causative. Similar as to the adjusted full model, breastfeeding showed no association with asthma (Table 4, see also Figure 3A).

**Table 4.** Results from meta-analyzing the matched groups for the fixed-effect model (P and OR) and for the random-effect model (P(R) and OR(R)). Q is the p-value for the Cochran’s Q statistics, I is the I^2 heterogeneity index (0-100) and N is the number of included groups. Each group is defined in Figure 1.

| Disease | P | P(R) | OR | OR(P) | Q | I | N |
| --- | --- | --- | --- | --- | --- | --- | --- |
| Asthma | 0.656 | 0.684 | 0.992 | 0.990 | 0.114 | 31.1 | 16 |
| Hay fever / Eczema | 0.00012 | 0.0022 | 1.052 | 1.048 | 0.2485 | 17.92 | 16 |
| Hay fever | $6.23 \times 10^{-5}$ | $6.23 \times 10^{-5}$ | 1.112 | 1.112 | 0.807 | 0.0 | 16 |
| Eczema | 0.0088 | 0.0088 | 1.105 | 1.105 | 0.792 | 0.0 | 16 |

## Discussion

We have studied the effect of breastfeeding on risk of being diagnosed with atopic disease in a population-based cohort from the UK, with participants born between 1939 and 1970. Frequency of breastfeeding decreased from over 80% in participants born between 1937 and 1941, to below 50% in participants born in 1970. This trend was probably due to the emergence of milk substitutes on the market in the 1950s, which potentially influenced breastfeeding practices. Worldwide, breastfeeding reached an all-time low in 1971, with as few as 27% of mothers from the United States breastfeeding their babies^15^. During the same time period (1939 to 1970), we can see an increase in the incidence of asthma, hay fever and eczema in the cohort (Figure 1). Therefore, not adjusting for year of birth results in a strong negative association between disease risk and breastfeeding. However, after including year of birth in the logistic un-matched model the effect of breastfeeding changed from protective to conferring increased odds for hay fever and eczema. These results show how important it is to adjust for appropriate confounders.

Identifying the actual effect of breastfeeding on disease outcome is a challenge since there may be ignored or unknown and unobserved differences between breastfed and non-breastfed participants. For example, non-linear effects and interactions between covariates could mask or distort the effect of breastfeeding. Our initial approach was to linearly correct for all associated covariates. Even after adjusting for possible confounders, it is still possible to that effects might reflect differences between the breastfed and the non-breastfed group. The best method to overcome such problems would be a randomized trial, where mothers are randomized into breastfeeding and or not and where the babies are followed up for a lifetime. However, this is not a feasible approach as breastfeeding is a critical component of child rearing. Another way to overcome such problems, and to strengthen the results from our main analysis, is by matching breastfed and non-breastfed participants for suspected confounders. After matching breastfed and non-breastfed participants, the results where still significant with the same direction of effect, which strengthens our conclusion that breastfeeding is associated with an increased odds of being diagnosed with hay fever and eczema (Table 4, Supplementary Table 2, Figure 3B-D), but that there is no detectable effect of breastfeeding on asthma.

It is remarkable and quite unexpected that we see an association between breastfeeding and an increased risk for hay fever and eczema. One hypothesis for such effect can be contaminants present in breast milk. Previous studies have identified increased level of contaminants in breast milk^16,17,18^, which agrees with this finding. Both hay fever and eczema are also heritable^19^ and it is possible that mothers with allergies more often breastfed their babies compared to mothers without any allergy. It is also important to remember that we might detect a reverse causality, that individuals with a known risk of developing atopic diseases (for example participants with atopic disease within their family) might not have been breastfed or had a higher chance of being breastfed due to recommendations.

We could also see an effect of socioeconomic status (for which Townsend deprivation index is used as a proxy) on asthma, hay fever and eczema. The Townsend deprivation index is a measurement of neighborhood-level of unemployment, non-car ownership, non-home ownership, and household overcrowding across UK. High socioeconomic status was associated with lower odds for asthma but higher risk for hay fever and eczema. The increased odds for hay fever and eczema is in line with the western world hygiene hypothesis, which states that growing up in a cleaner environment increases the risk of being diagnosed with hay fever and eczema due to a lack of early childhood exposure to for example, microorganisms^20^.

Maternal smoking around birth and smoking in house seem to be risk factors for developing asthma, while they have a protective effect on being diagnosed with hay fever and eczema. Maternal smoking around birth and being exposed to smoke have shown conflicting results in previous studies, some studies reporting an increased risk and some reporting a protective effect against atopic sensitization^21,22,23^. A previous study reported a protective effect of maternal smoking, even after correcting for maternal disease incidence^24^. This protective effect might be similar to the hygiene hypothesis^20^. However, even though this study showed a protective effect of maternal smoking around birth and being exposed to tobacco smoke at home on the risk of being diagnosed with hay fever and eczema, there are a number of important reasons not to expose other people to tobacco smoke, for example the fact that second hand smoke increases the risk of poor lung development^25^, lung cancer^26^ and heart disease^27^.

Previous studies have reported conflicting results, which might be due to random false positives, heterogeneity between studies and depending on the confounders that were considered. This study has a great power, compared to previous studies, due to the large sample size and homogeneous population. We also considered a large number of known confounders and matched breastfed and non-breastfed for important confounders. However, there were also a number of limitations in this study that we will discuss.

### Limitation

Due to the dichotomous nature of the data, we could not investigate the effect of duration of breastfeeding (more than 2 month), a dose-response effect (2,4, or 6 month), or the effect of exclusive breastfeeding. Both duration and exclusivity of breastfeeding are important since dose-response is well established for many breast milk benefits, especially for child mortality^28,29^. However, in previous studies that investigated the association between breastfeeding and atopic diseases, it appears as dose-response effect and duration do not change the direction of effect, rather only increases the effect size, which, in turn, influence the size of the P-values^8,12,6^. Lacking this information will therefore most likely only decrease our power to find significant associations, and not pick up effects that are false positives or that have opposite effects, as we detected when not including year born in this study. Furthermore, we could not adjust for family history of disease, which might introduce some bias. However, a previous study showed that breastfeeding effects are not affected by parental history of hay fever or asthma^12^. We can’t make any statements on possible different effects of breastfeeding between different risk groups depending on the level of prevalence, or lack of disease in families.

Even though the phenotypes are self-reported in the UK Biobank database, which may lead to an overestimation of disease incidence, the questions are well defined and identical for all participants. Using self-reported data from such a large population based cohort as UK Biobank, the need for a strict diagnosis criteria is less important due to the large power.

We adjusted for TDI, which is a proxy for socioeconomic status of the person being breastfed, not for the home they grow up in. Since we see the same pattern as what has previously already been established scientifically, i.e. that lower socioeconomic status increase the risk of asthma and lowering the risk of hay fever, we don’t believe that this limitation will give a large bias to the analysis^18,30,31^.

Understanding the limitations of this study should not limit the value of it, which is an important contribution to the breastfeeding and risk prevention of atopic disease literature.

## Conclusion

This study reports evidence that breastfeeding is associated with an increased risk for hay fever and eczema. Due to the high power, achieved by our large sample size and rigorous information on potentially confounding variable, we can conclude that breastfeeding is not likely to have a large effect on risk of asthma. The study also show that being exposed to smoke around birth increase the risk of asthma while lowering the risk of being diagnosed with hay fever and eczema. These results should not be used to recommend breastfeeding or to discourage it since the present study only investigates the association between breastfeeding history and being diagnosed with asthma, hay fever and eczema during lifetime. Therefore, the authors of the present study do not take any positions as to whether breastfeeding represents the best source of nutrition for newborns.

## Acknowledgment

This research was conducted using the UK Biobank Resource under Application Number 15479. The computations were performed on resources provided by SNIC through Uppsala Multidisciplinary Center for Advanced Computational Science (UPPMAX) under projects b2016021203.

## Author’s Contributions

Planned the study (WEE and ÅJ), analyzed the data (WE, TK and ÅJ), literature search (WEE), Figures (WEE), data interpretation (WEE, ÅJ, TK, MRA, CAH), writing of manuscript (WEE, ÅJ, TK, MRA, CAH).

